# A Single Lipid Nanoparticle Formulation Enables Delivery of Diverse RNA Therapeutics to Human Muscle Models of Duchenne Muscular Dystrophy

**DOI:** 10.64898/2026.07.29.741396

**Authors:** Paola Galbiati, Delphine Leclerc, Margaux Mombled, Ruqayya Khan, Maëlle Ralu, Giorgia Bimbi, Giulia Scalisi, Kamel Mamchaoui, Francesco Saverio Tedesco, Sonia Albini, Mario Amendola

## Abstract

Duchenne muscular dystrophy is a lethal neuromuscular disorder caused by the absence of dystrophin, for which no curative treatment is available. RNA-based approaches have shown promising results; however, their evaluation is hindered by the lack of robust and rapid delivery methods for differentiated human muscle cells, which represent the most physiologically relevant *in vitro* models for assessing therapeutic strategies.

Here, we establish a versatile lipid nanoparticle platform enabling efficient delivery of diverse RNA therapeutics across a range of human muscle models, including myotubes, induced pluripotent stem cell-derived myotubes, myoblasts, cardiomyocytes, and 3D engineered skeletal muscle tissues. Remarkably, a single commercially available lipid nanoparticle formulation supports delivery of cargos spanning more than 300-fold in size, from short antisense oligonucleotides (∼20 nt) to complex CRISPR-based editors (up to ∼6.7 kb), including Cas9 nucleases, adenine base editors, and CRISPRa systems. This enables efficient gene correction and transcriptional modulation, resulting in dystrophin restoration or compensatory utrophin upregulation in relevant Duchenne muscular dystrophy models. Together, our results establish a single lipid nanoparticle formulation as a versatile platform for RNA delivery in human muscle systems and provide a practical framework for the rapid preclinical assessment of emerging therapies for Duchenne muscular dystrophy and other neuromuscular disorders.

**GRAPHICAL ABSTRACT:** 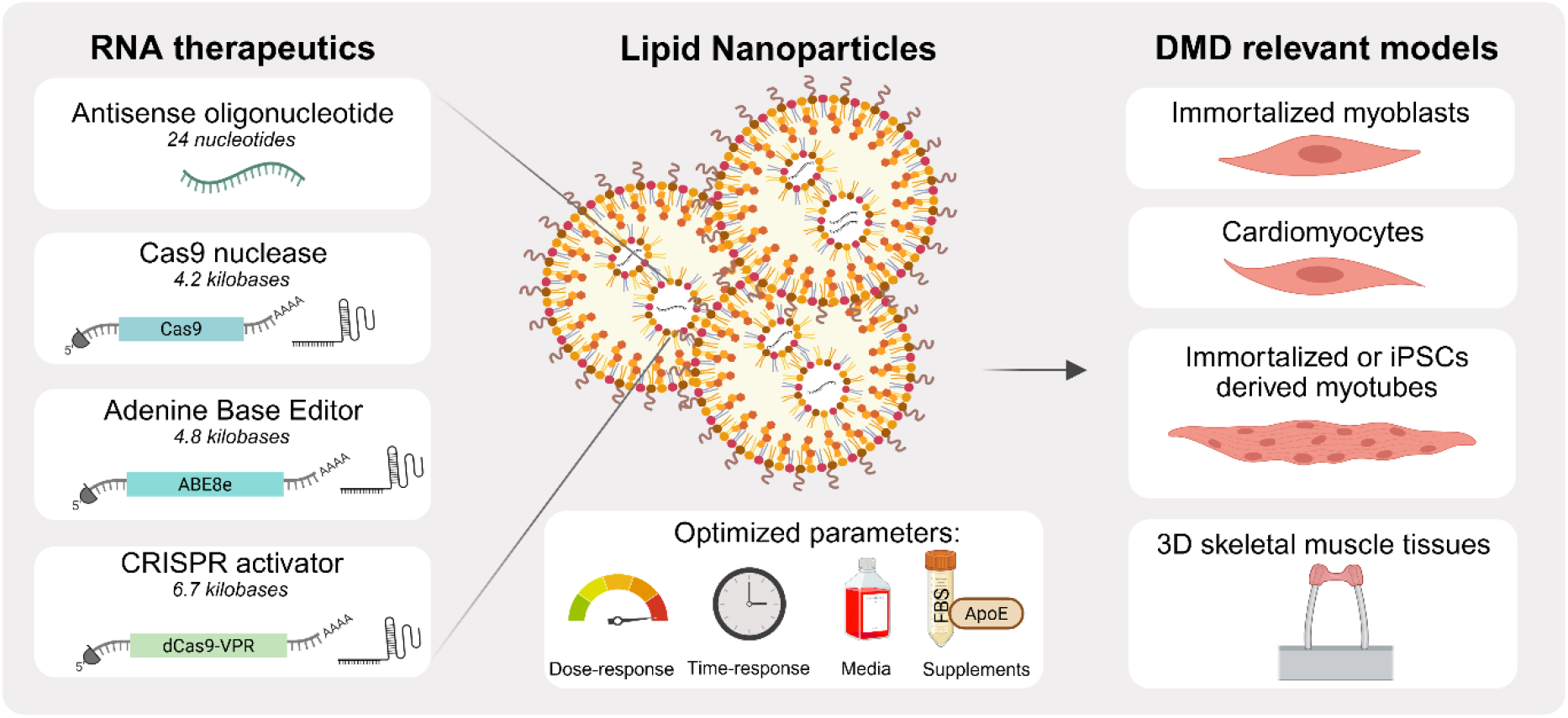

*Created in BioRender. Leclerc, D. (2026)* https://BioRender.com/90h0fme

## INTRODUCTION

Duchenne muscular dystrophy (DMD) is a severe X-linked neuromuscular disorder affecting approximately 1 in 5,000 newborn males. It is caused by pathogenic variants in the *DMD* gene that abolish the expression of dystrophin, a crucial structural protein required to maintain muscle fiber integrity during contraction. The large size of the *DMD* gene (the largest in nature: ∼2.4 Mb, 79 exons) results in extensive mutational heterogeneity, with thousands of distinct pathogenic variants identified in patients [1]. Clinically, DMD is characterized by progressive muscle degeneration leading to loss of ambulation around 12 years of age, followed by severe respiratory and cardiac complications that ultimately result in premature death in early adulthood [1]. Despite major therapeutic advances, no effective curative therapy is currently available.

The most of ongoing therapeutic strategies aim to restore dystrophin expression. While Adeno associated viral vector (AAV) mediated microdystrophin delivery has shown clinical promise, its application remains limited by restricted cargo capacity, incomplete functional rescue, pre-existing immunity, manufacturing constraints, and recent safety concerns, including fatal immune reactions [2,3]. Alternative RNA-based approaches, including splice-switching antisense oligonucleotides (ASOs) [4,5], and CRISPR-based genome-editing agents [6,7], have emerged as promising alternatives for mutation-specific correction strategies. However, their development is constrained by the lack of efficient, scalable, and standardized delivery methods in relevant human muscle models.

Current human muscle models, including proliferative myoblasts, post-mitotic multinucleated myotubes, cardiomyocytes and three-dimensional (3D) organoids, provide physiologically relevant platforms for evaluating therapeutic strategies [8,9]. Among these, differentiated myotubes represent one of the most relevant yet technically challenging platforms for evaluating RNA therapeutics aimed at restoring dystrophin expression. Unlike proliferating myoblasts, myotubes robustly express key muscle-specific proteins, including dystrophin [10]. Furthermore, genome-editing outcomes are dependent on cell-cycle stage, with non-dividing cells (myotubes) relying on DNA repair mechanisms that differ markedly from those active in proliferating myoblasts [11]. Consequently, direct evaluation of RNA therapeutics in differentiated muscle cells is essential for accurately predicting therapeutic performance.

Despite their relevance, efficient nucleic acid delivery into differentiated muscle cells remains a major technical challenge. Conventional transfection and electroporation approaches perform poorly with substantial toxicity in myotubes due to their fragile and multinucleated nature. Although recent studies have optimized lipoplex-based transfection reagents for use in muscle cells, efficient and reproducible delivery to post-mitotic myotubes remains a major unresolved challenge [12]. As a result, many studies perform genetic modification in proliferating myoblasts before inducing differentiation, a workflow that may not faithfully reflect delivery efficiency, gene-editing outcomes, or therapeutic responses in mature muscle tissue [13,14].

Viral vectors such as AAVs [15,16] and lentiviral vectors [17] can improve delivery efficiency, but they introduce additional challenges, including production complexity, limited cargo size, and limited suitability for transient genome-editing applications. Altogether, these limitations highlight the need for a versatile, efficient, and scalable RNA delivery platform applicable across diverse human muscle systems. Lipid nanoparticles (LNPs) have emerged as versatile non-viral delivery vehicles capable of encapsulating a wide range of nucleic acid, protein and drug cargos. LNPs are typically composed of four key components: ionizable lipids, PEGylated lipids, phospholipids, and cholesterol. Each component plays a critical role in nanoparticle stability, cargo encapsulation, intracellular delivery, and overall safety profile. Importantly, variations in LNP composition can substantially influence biodistribution, delivery efficiency, and cell-type specificity [18]. Cellular uptake of LNPs primarily occurs through receptor-mediated endocytosis, notably involving receptors such as the low-density lipoprotein receptor [19].

Following internalization, endosomal acidification protonates the ionizable lipid, promoting endosomal membrane destabilization and facilitating cargo release into the cytoplasm.

In addition to their favorable delivery properties, LNPs are cost-effective, reproducible, compatible with scalable microfluidic manufacturing, and have already demonstrated clinical utility in multiple contexts [20–22]. Importantly, LNP-mediated RNA delivery enables transient expression of therapeutic cargos, allowing hit-and-run expression of genome-editing tools, which reduces prolonged nuclease exposure and associated genotoxic risks. Despite these advantages, the potential of LNPs for RNA delivery in differentiated myotubes and advanced muscle models remains relatively unexplored [12,23]

Here, we assessed a single commercially available LNP formulation as a reference platform, chosen for its efficient RNA encapsulation and delivery efficiency, low toxicity across diverse cell types, and compatibility with scalable microfluidic manufacturing. We first establish efficient RNA delivery in differentiated human myotubes, including immortalized and iPSC-derived models, before extending this approach to myoblasts, cardiomyocytes and engineered 3D skeletal muscle tissues. We further demonstrated efficient delivery of diverse RNA cargos spanning more than 300-fold in size, from short ASOs (∼20 nt) to large CRISPR-based effectors (∼6.7 kb), without the need for protocol re-optimization. This work provides a single platform for rapid and scalable preclinical evaluation of RNA-based therapies for DMD and related neuromuscular disorders.

## RESULTS

### LNPs overcome transfection barriers in human post-mitotic myotubes

Efficient delivery of nucleic acids into differentiated skeletal muscle cells remains a major bottleneck for the evaluation of RNA-based therapeutics. Although several approaches have been optimized for myogenic cells, multinucleated post-mitotic myotubes remain particularly challenging to transfect and are therefore underutilized in therapeutic development pipelines [24].

To address this challenge, we first compared plasmid DNA and mRNA nucleofection of GFP in human immortalized myoblasts derived from a Duchenne muscular dystrophy (DMD) patient with a deletion between exons 45-50 (ΔEx45–50). Consistent with previous reports [25], plasmid DNA nucleofection resulted in low and heterogeneous GFP expression in myogenic cells, while mRNA supported robust, homogeneous, and dose-dependent transgene expression without affecting cell viability (Fig. S1A-D). Based on these results, mRNA was selected as the preferred cargo for the subsequent experiments.

We then evaluated if LNP could be used to deliver mRNA in differentiated human myotubes derived from immortalized wild-type and Duchenne muscular dystrophy (DMD) myoblasts (Fig. 1A). We generated LNP loaded with an mRNA coding for GFP reporter and controlled encapsulation efficiency (∼96%) and particle diameter and homogeneity (∼96 nm) (Fig. S2A-C). Given that LNP uptake is predominantly mediated by the low-density lipoprotein receptor (LDLr), we first confirmed LDLr expression in human wild-type and DMD myotubes by RT-qPCR analysis (Fig. S3A). Comparison of different culture conditions revealed that LNP transfection performed in growth medium markedly increased GFP expression compared with differentiation medium, either alone or supplemented with ApoE3, ApoE4 or serum (Fig. 1B). Using these optimized conditions, LNPs enabled robust and dose-dependent increase in fluorescence intensity in both wild-type and DMD myotubes, without apparent toxicity (Fig. 1C-D, S3C-D). Importantly, LNP-mediated delivery significantly outperformed Lipofectamine (Fig. S3B), establishing LNPs as a superior delivery modality in differentiated muscle cells.

**Figure 1.**
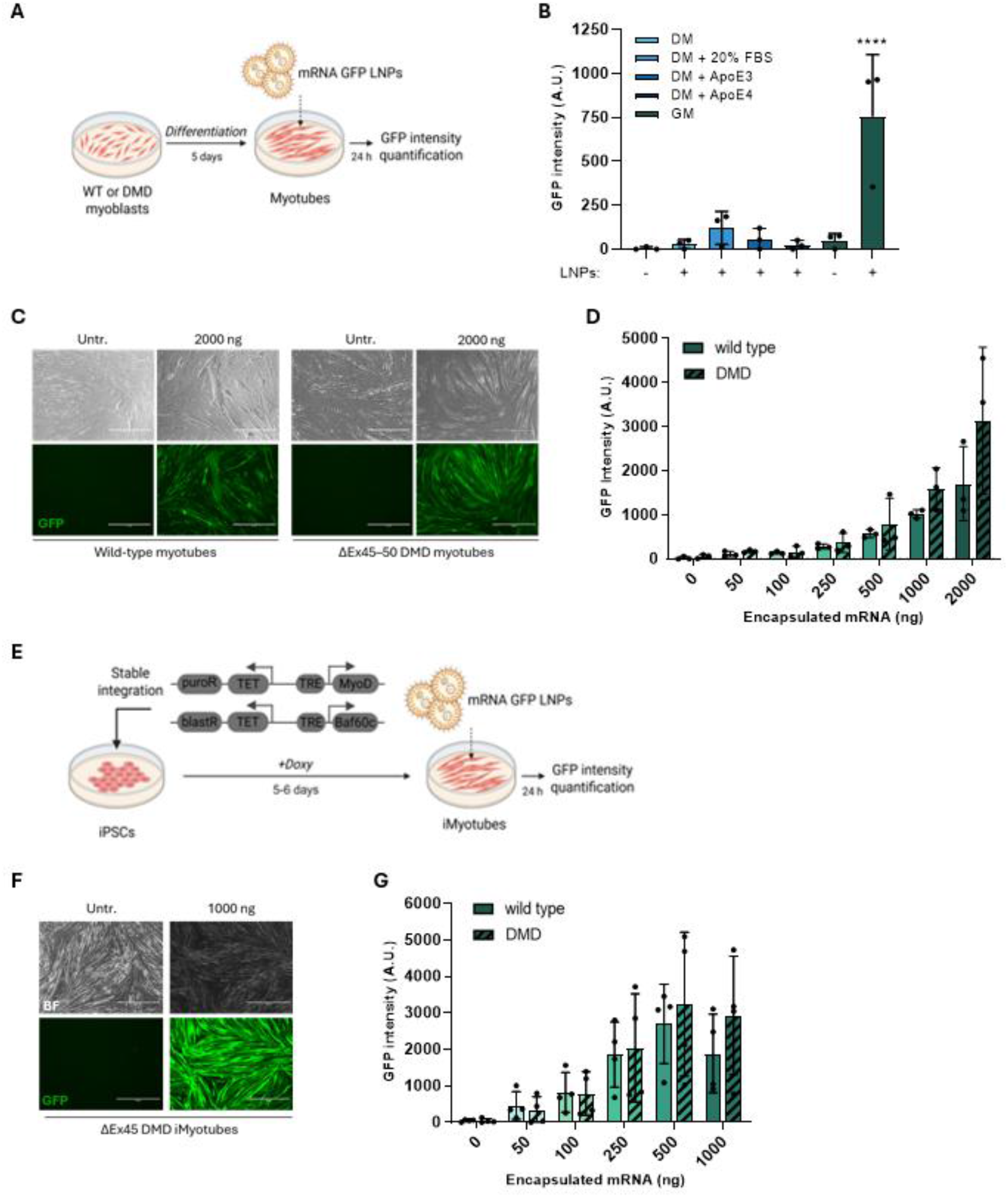
LNPs can efficiently deliver GFP mRNA to human immortalized and iPSC-derived myotubes from unaffected individuals and DMD patients. (**A**) Schematic representation of the experimental workflow for immortalized myotubes transfection. (**B**) GFP intensity quantification in ΔEx45–50 DMD myotubes transfected with 500 ng of LNP-GFP mRNA. Transfection was performed in differentiation medium (DM) - with or without ApoE3, ApoE4, or 20% FBS - and compared with delivery in growth medium (GM). (**C**) Representative fluorescence images of wild-type and ΔEx45–50 DMD myotubes untransfected or transfected with 2000 ng of LNP-GFP mRNA. Scale bar: 400 μm. BF = brightfield. (**D**) GFP intensity quantification in wild-type and ΔEx45–50 DMD myotubes transfected with increasing doses of LNP-GFP mRNA. (**E**) Schematic representation of the experimental workflow for iPSCs differentiation and iMyotubes transfection. (**F**) Representative fluorescence images of ΔEx45 DMD iMyotubes untransfected or transfected with 1000 ng of LNP-GFP mRNA. Scale bar: 1000 μm. BF = brightfield. (**G**) GFP fluorescence intensity in wild-type and ΔEx45 DMD iMyotubes transfected with increasing doses of LNP-GFP mRNA. Statistical analyses were performed using a two-tailed Student’s *t*-test for single comparisons and one-way ANOVA followed by Dunnett’s multiple-comparisons test for multiple comparisons. **** p < 0.0001. Each dot represents one sample.

To determine whether these findings could be extended to more physiologically relevant human muscle models, we next evaluated LNP-mediated delivery in primary human myotubes derived from human induced pluripotent stem cells (iMyotubes) generated from a healthy control and a DMD patient carrying an exon 45 deletion (ΔEx45) (Fig. 1E). Following confirmation of LDLr expression (Fig. S3E), LNP-mediated delivery resulted in robust and dose-dependent GFP expression, reaching nearly 100% GFP-positive cells (Fig. 1F-G, S3F).

Together, these findings establish LNPs as an efficient platform for RNA delivery into differentiated human skeletal muscle cells, including both immortalized and iPSC-derived myotubes.

### A single LNP formulation enables efficient RNA delivery across diverse human muscle models

Having established efficient delivery in differentiated muscle cells, we investigated whether the same LNP formulation could support efficient RNA delivery across multiple human muscle systems without protocol re-optimization.

First, we evaluated LNP-GFP mRNA delivery in proliferating immortalized myoblasts derived from a healthy donor and a DMD patient (Fig. 2A). We confirmed LDLr expression (Fig. S4A) and efficient LNP transfection in both cell types, with a clear dose-dependent and time-dependent increase in GFP-positive cells and fluorescence intensity, without affecting cell viability, again demonstrating superior performance compared with Lipofectamine-mediated delivery (Fig. 2B-D, Fig. S4B-G). Transfection efficiencies were comparable to those obtained by direct mRNA nucleofection while avoiding the complexity associated with electroporation-based approaches.

**Figure 2.**
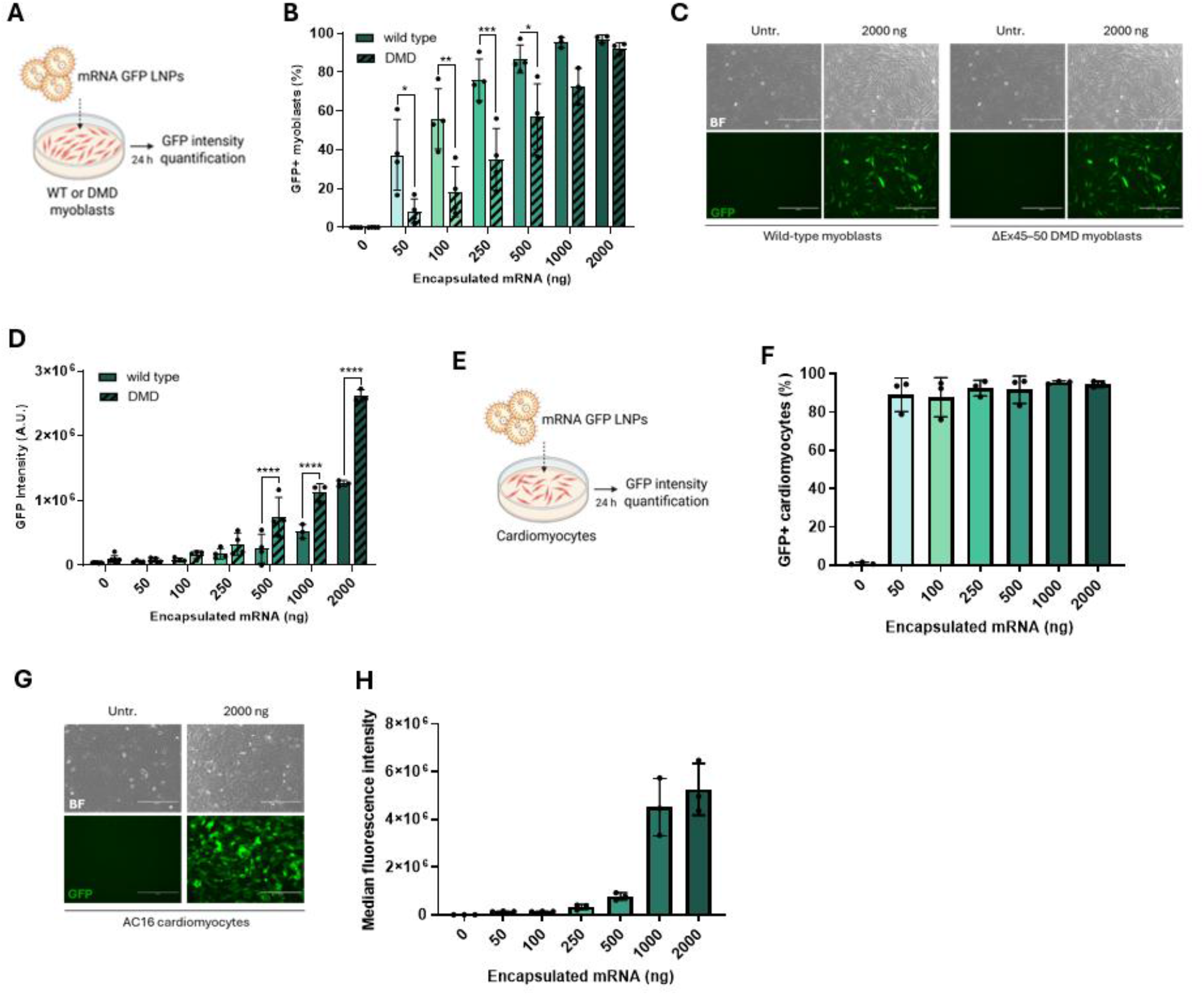
LNPs can efficiently deliver GFP mRNA to human immortalized myoblasts from unaffected individuals and DMD patients, and AC16 cardiomyocytes. (**A**) Schematic representation of the experimental workflow for immortalized myoblasts transfection. (**B**) Flow-cytometry quantification of GFP-positive cells of wild-type and ΔEx45–50 DMD myoblasts transfected with increasing doses of encapsulated GFP mRNA. (**C**) Representative fluorescence images of ΔEx45–50 DMD myoblasts untrasfected or transfected with 2000 ng LNP–GFP mRNA across a range of doses. Scale bar: 400 μm. BF = brightfield. (**D**) Flow-cytometry quantification of GFP fluorescence intensity of wild-type and ΔEx45–50 DMD myoblasts transfected with increasing doses of encapsulated GFP mRNA. (**E**) Schematic representation of the experimental workflow for AC16 cardiomyocytes transfection. (**F**) Percentage of GFP-positive AC16 cardiomyocytes following transfection with GFP mRNA at increasing doses. **(G)** Representative fluorescence images of AC16 cardiomyocytes untrasfected or transfected with 2000 ng LNP–GFP mRNA across a range of doses. Scale bar: 400 μm. BF = brightfield. (**H**) Cytometry-based quantification of median GFP fluorescence intensity of AC16 cardiomyocytes transfected with GFP-encoding mRNA at increasing doses. Statistical analyses were performed using a two tailed Student’s t test for single comparisons and one way ANOVA followed by Dunnett’s multiple comparisons test for multiple comparisons. NS, not significant, * p < 0.05, ** p < 0.01, *** p < 0.001, **** p < 0.0001. Each dot represents one sample.

Because cardiac dysfunction represents a major contributor to morbidity and mortality in DMD patients, we also tested LNP-GFP delivery in AC16 human cardiomyocytes (Fig. 2E). LDLr expression was readily detected in these cells (Fig. S4H) and, similarly to myoblasts and myotubes, we observed a dose dependent increase in GFP expression, with ∼100% GFP-positive cells detected at all tested doses (Fig. 2F-H, S4I). To determine whether LNP-mediated delivery extends beyond conventional monolayers (2D), we evaluated their performance in three-dimensional (3D) engineered skeletal muscle tissues, which better model the complex architecture of muscle tissue. When LNP-GFP were incorporated into the cell–hydrogel mixture during casting, we observed uniform and robust GFP expression throughout the whole organoid, as assessed by whole-mount fluorescence and confocal microscopy (Fig. 3A-B). We then evaluated delivery in pre-formed mature tissues, where GFP expression remained detectable, although more localized to peripheral regions, consistent with limited nanoparticle diffusion (Fig. 3C). Collectively, these results demonstrate that a single commercially available LNP formulation supports efficient RNA delivery across a broad spectrum of DMD-relevant cellular models, including post-mitotic myotubes, proliferating skeletal muscle cells, and cardiomyocytes as well as 3D skeletal muscle models.

**Figure 3.**
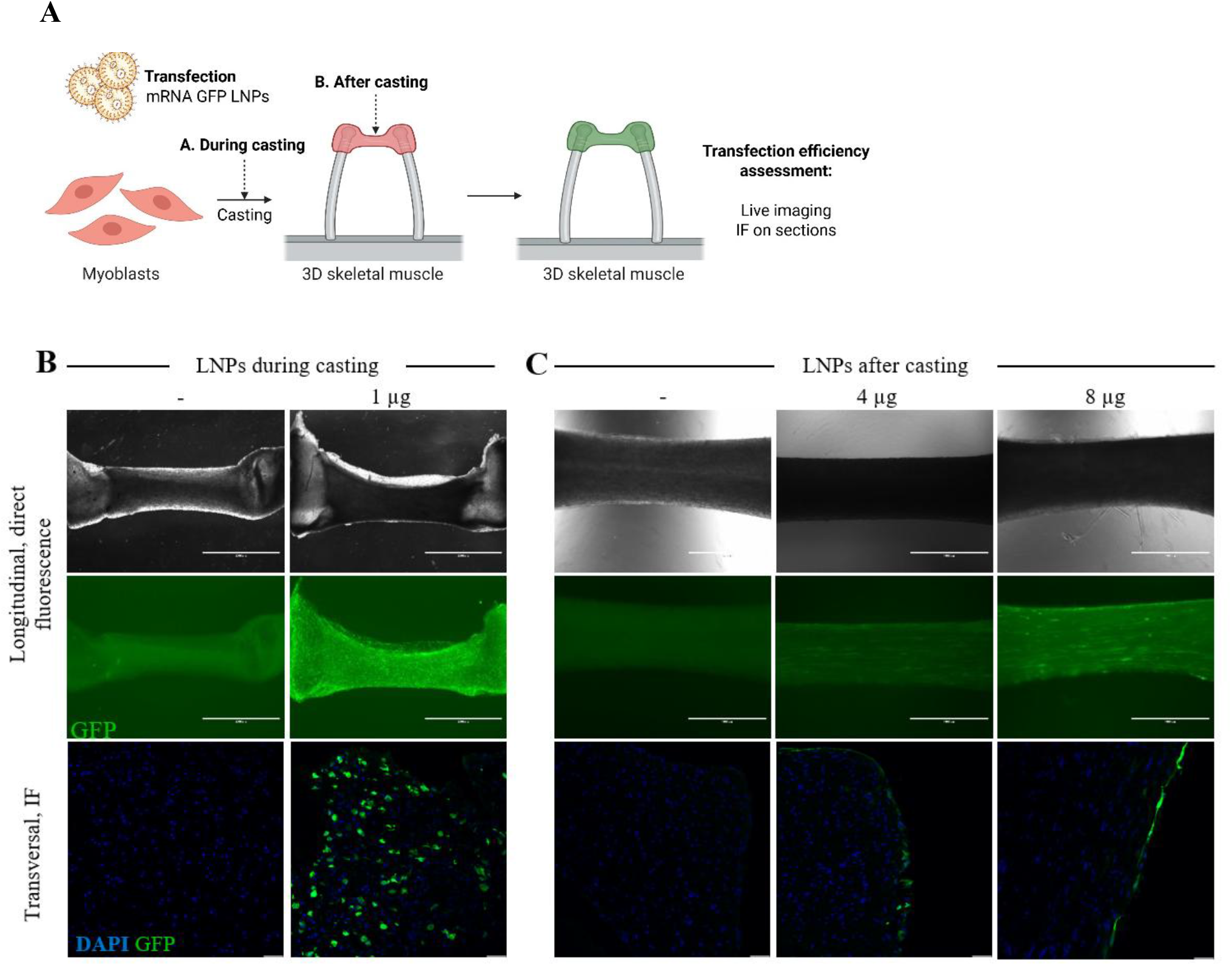
LNPs enable mRNA delivery in 3D skeletal muscle tissues. (**A**) Schematic representation of the experimental workflow. (**B-C**) 3D skeletal muscles were transfected (B) during casting with 1 µg or after casting with 4 µg or 8 µg of LNP-GFP mRNA and analyzed 24 h after. Top row: Brightfield images of longitudinal sections (scale bar: 2000 µm); Middle row: direct GFP fluorescence of longitudinal sections. Bottom row: GFP immunofluorescence of transversal sections; nuclei are stained with DAPI (scale bar: 50 µm).

### LNP enables efficient delivery of therapeutic ASO in post-mitotic myotubes

Having established efficient RNA delivery across multiple DMD-relevant cellular models, we next evaluated whether LNPs could efficiently deliver therapeutic RNA cargos.

To this end, we encapsulated a splice-switching antisense oligonucleotide (ASO, 24 nt) targeting exon 44 of the *DMD* gene and applied it to DMD iMyotubes carrying an exon 45 deletion [9] (Fig. 4A) (Brogidirsen, NCT05135663) [26]. LNP-mediated ASO delivery resulted in dose-dependent exon 44 skipping at the mRNA level, reaching up to ∼45% (Fig. 4B). This corrected mRNA was validated by Sanger sequencing confirming the expected exon 43–exon 46 splice junction (Fig. 4C) and resulted into dose-dependent restoration of dystrophin protein expression (Fig. 4D).

**Figure 4.**
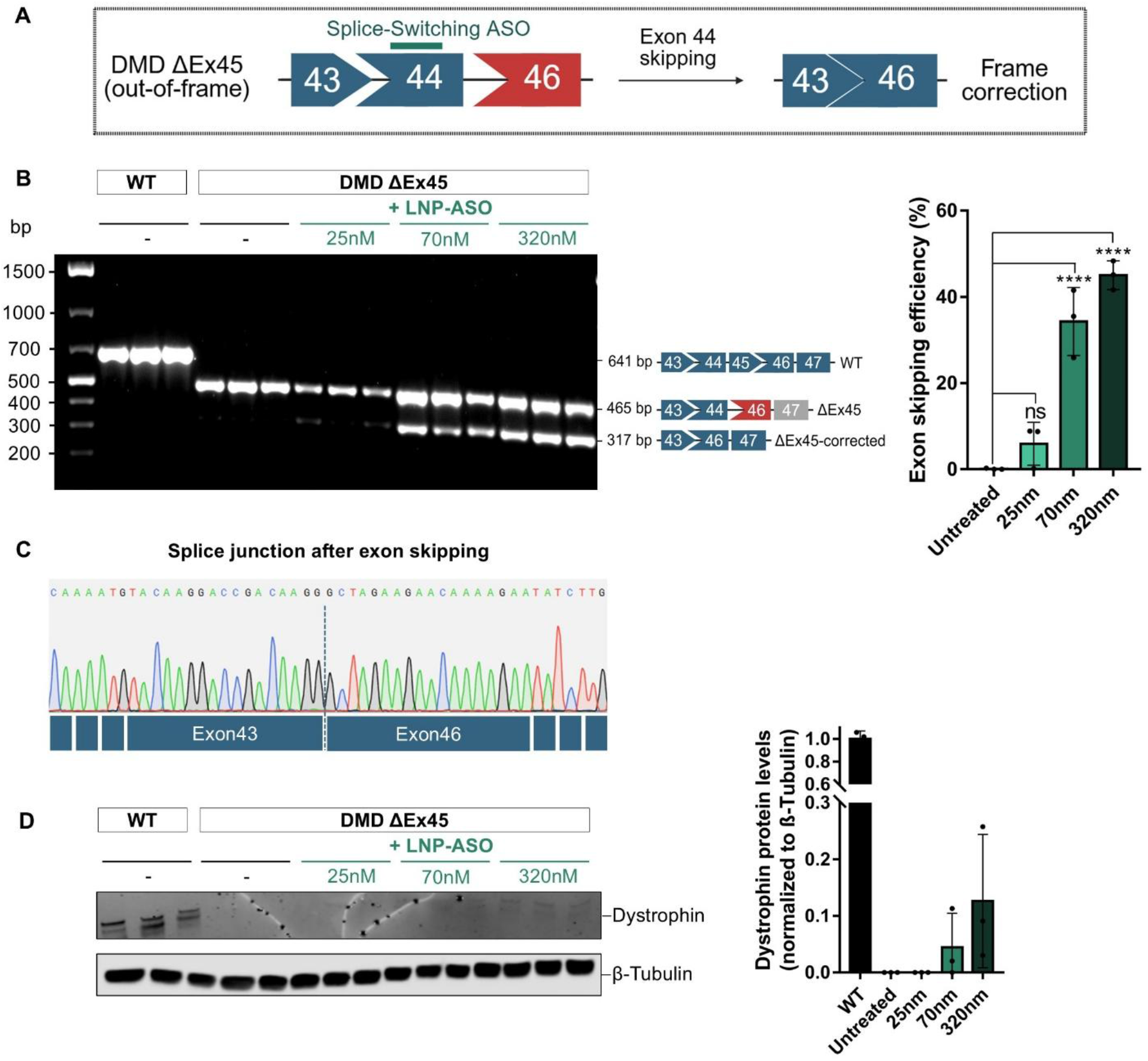
Efficient LNP-ASO delivery induces DMD exon 44 skipping in DMD ΔEx45 iMyotubes. (**A**) Schematic representation of the exon skipping strategy. (**B**) RT-PCR analysis shows exon 44 skipping after LNP-ASO treatment. Quantification of exon-skipping efficiency is shown on the right. (**C**) Exon skipping was confirmed by Sanger sequencing, validating the new in-frame splice junction between DMD exons 43 and 46. (**D**) Western blot analysis of dystrophin expression, with β-tubulin used as a loading control. Quantification of dystrophin levels normalized to β-tubulin is shown on the right and expressed relative to wild-type levels. Data represent mean ± S.D. (n = 3 independent biological replicates). Statistical significance was determined using a one-way ANOVA followed by Dunnett’s multiple comparisons test relative to the control. NS, not significant, **** p < 0.0001. Each dot represents one sample.

These findings demonstrate that LNPs efficiently deliver small, chemically modified RNA therapeutics and promote molecular correction in post-mitotic human muscle cells, extending beyond simple transgene expression.

### LNP enables efficient delivery of different CRISPR/Cas9 editor mRNAs in post-mitotic human myotubes and cardiomyocytes

The transient nature of mRNA delivery is well suited for the hit-and-run expression of genome-editing agents; therefore, we decided to test if LNPs could deliver both the large and complex Cas9-derived mRNAs (∼4.2 kb to ∼6.7kb) together with their sgRNA cargos. Interestingly, the distribution and average LNP sizes did not change with the size of the RNA cargo (Fig. S2A-C).

LNPs loaded with Cas9 mRNA and a sgRNA targeting *DMD* exon 53 were delivered to immortalized DMD myotubes carrying an exon 52 deletion [27] (Fig. 5A). The sgRNA was designed just upstream of the premature stop codon in exon 53, to promote reframing through Non-Homologous End-Joining mediated +1 insertions following Cas9 cleavage (Fig. 5B). Three days post-transfection, Sanger sequencing analysis revealed efficient insertions/deletions (INDELs) formation, with a preference for reframing (+1) INDELs (Fig. 5C-D; Fig. S5A-B). These edits resulted in partial restoration of dystrophin protein expression (Fig. 5E, Fig. S5C).

**Figure 5.**
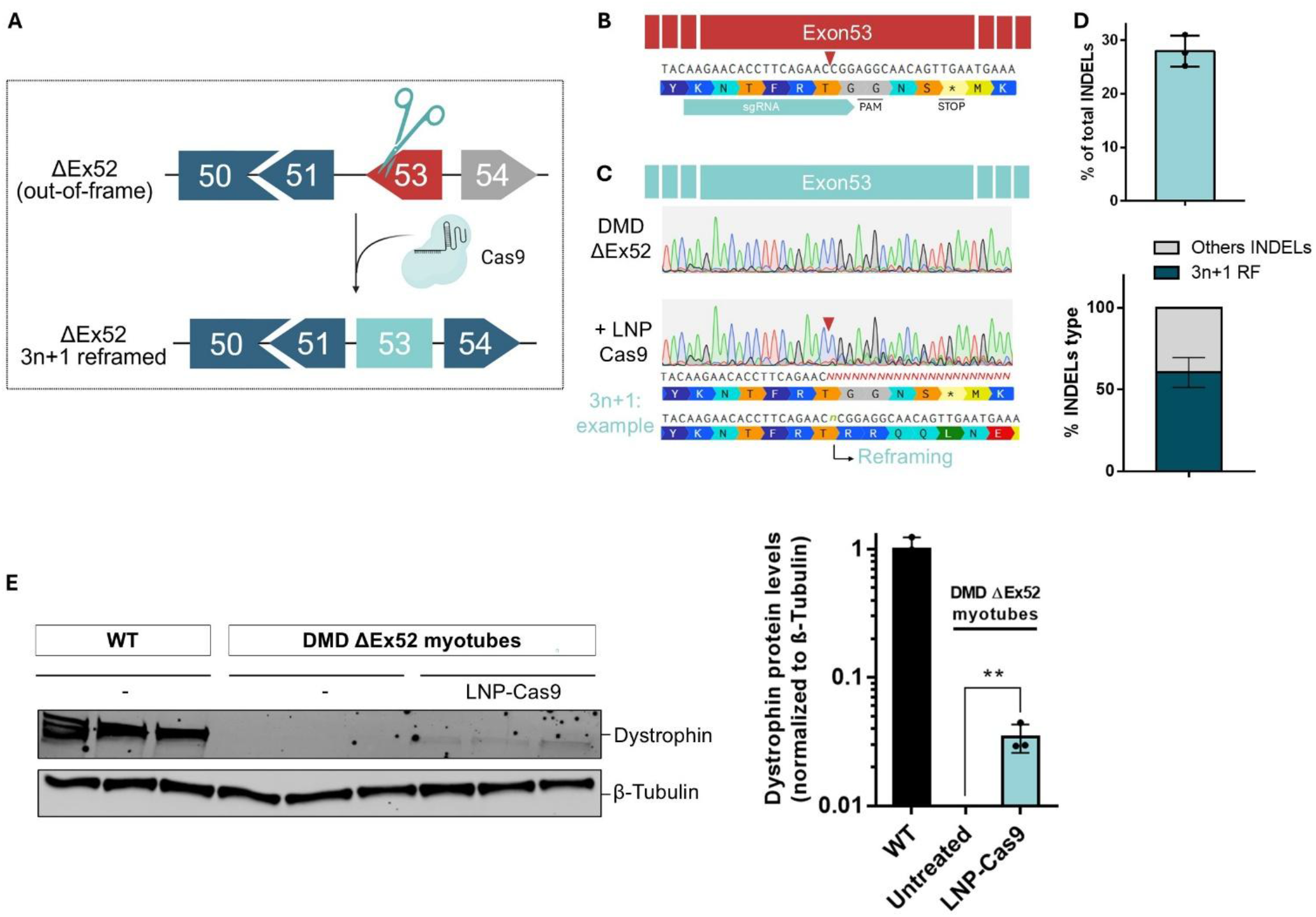
Efficient LNP-Cas9 delivery enables *DMD* reframing and dystrophin restoration in DMD ΔEx52 myotubes. (**A**) Schematic representation of the reframing strategy. (**B**) sgRNA binding site in dystrophin exon 53, located just upstream of the premature stop codon. The red arrowhead indicates the Cas9 cutting site. (**C**) Representative Sanger chromatograms of untreated and LNP-Cas9-treated myotubes. The red arrowhead indicates the Cas9 cutting site. (**D**) Quantification of INDELs using TIDE in myotubes. Top: percentage of total INDELs; bottom: percentage of reframing INDELs. (**E**) Western blot analysis of dystrophin expression, with β-tubulin used as a loading control. Quantification of dystrophin levels normalized to β-tubulin is shown on the right and expressed relative to wild-type levels. Data represent mean ± S.D. (n = 3 biological replicates). Statistical analysis was performed using a two-tailed Student’s t-test. ** p < 0.01. Each dot represents one sample.

To further evaluate the versatility of the platform, we tested LNP-mediated delivery of increasingly complex genome editing systems. LNPs encapsulating the adenine base editor ABE8e (∼4.8 kb) and its sgRNA were used to restore dystrophin expression in myotubes harbouring a deletion of exon 51 of the *DMD* gene, one of the most common single-exon deletion mutations. The ABE8e/sgRNA disrupts the splice donor site of exon 50 and promotes its skipping, thereby restoring the dystrophin reading frame (Fig. 6A-B) [28]. LNPs delivery resulted in high editing efficiency, reaching 75% A-to-G conversion (A7) (Fig. 6C-D, Fig. S6A), and led to robust and precise exon 50 skipping (∼93%) (Fig. 6E-F). These molecular edits translated into substantial dystrophin restoration, as confirmed by protein analysis (Fig. 6G-H, Fig. S6B). Interestingly, LNPs could be re-dosed to increase editing levels in myotubes (Fig. S7).

**Figure 6.**
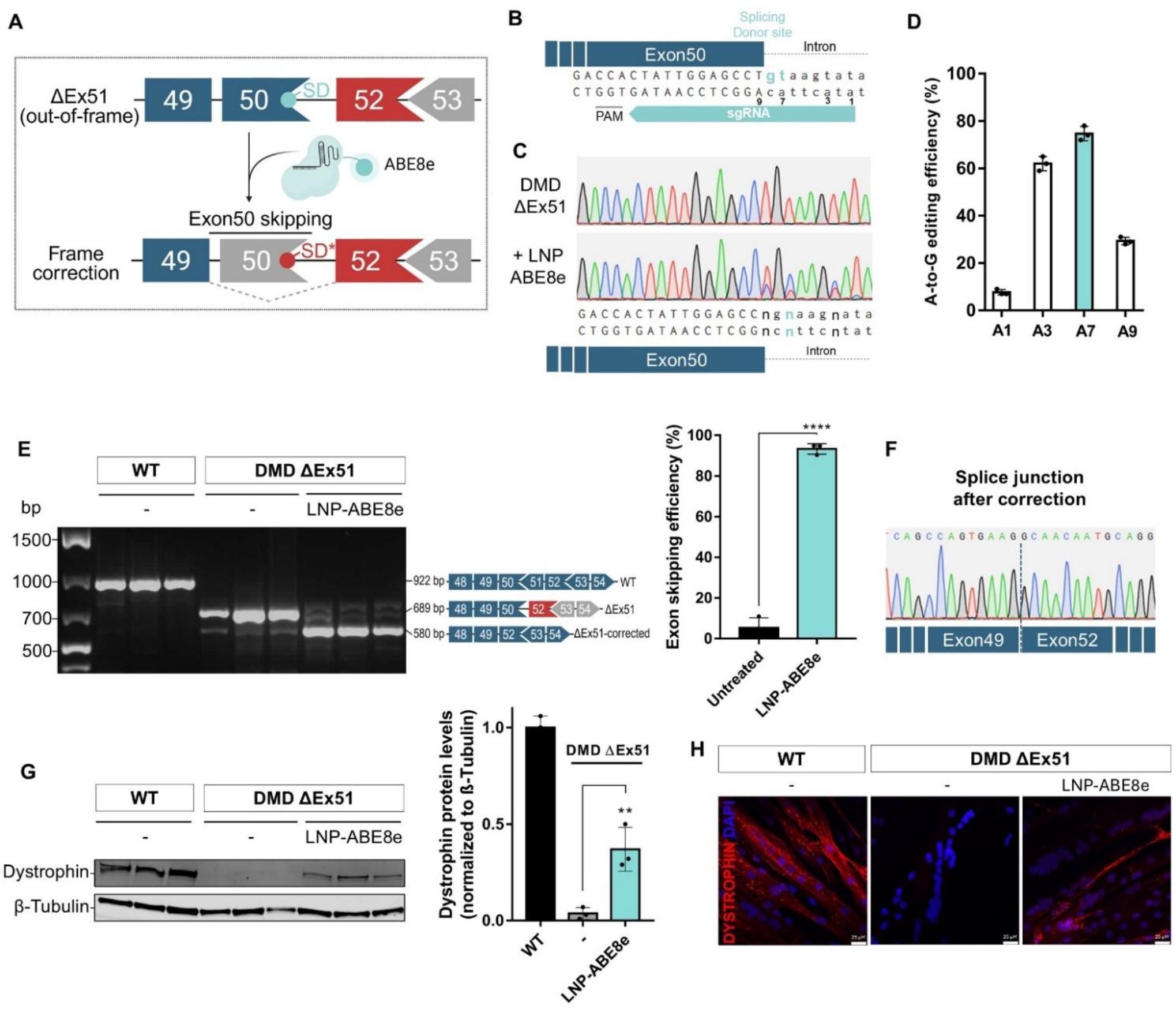
Efficient LNP-ABE8e delivery enables exon 50 skipping and dystrophin restoration in DMD ΔEx51 myotubes. (**A**) Schematic representation of the exon-skipping strategy. (**B**) sgRNA binding site in dystrophin exon 50 splice donor site; the targeted adenine (A7) within the ABE8e editing window is indicated. (**C**) Representative Sanger chromatograms of untreated and LNP-ABE8e–treated myotubes. (**D**) Quantification of A-to-G editing using EditR. (**E**) RT-PCR analysis shows exon 50 skipping after LNP-ABE8e treatment. Quantification of exon-skipping efficiency is shown on the right. (**F**) Exon skipping was confirmed by Sanger sequencing, validating the new in-frame splice junction between exons 49 and 52. (**G**) Western blot analysis of dystrophin expression, with β-tubulin loading control. Quantification of dystrophin levels normalized to β-tubulin is shown on the right and expressed relative to wild-type levels. (**H**) Immunofluorescence staining of dystrophin (red) and nuclei (DAPI, blue). Scale bar, 25 µm. Data represent mean ± S.D. (n = 3 biological replicates). Statistical analysis was performed using a two-tailed Student’s t-test. ** p < 0.01, **** p < 0.0001. Each dot represents one sample.

Lastly, we investigated whether LNPs could deliver even larger RNA cargos by testing a CRISPR activation system (dCas9-VPR, ∼6.7 kb) [29] targeting the endogenous utrophin (UTRN) promoter, a structural and functional paralogue of dystrophin, as a potential universal therapeutic strategy for DMD (Fig. 7A) [13,30,31]. LNP-mediated delivery resulted in robust upregulation of utrophin at both transcript and protein levels in ΔEx45–50 DMD myotubes and healthy cardiomyocytes (Fig. 7B–J).

**Figure 7.**
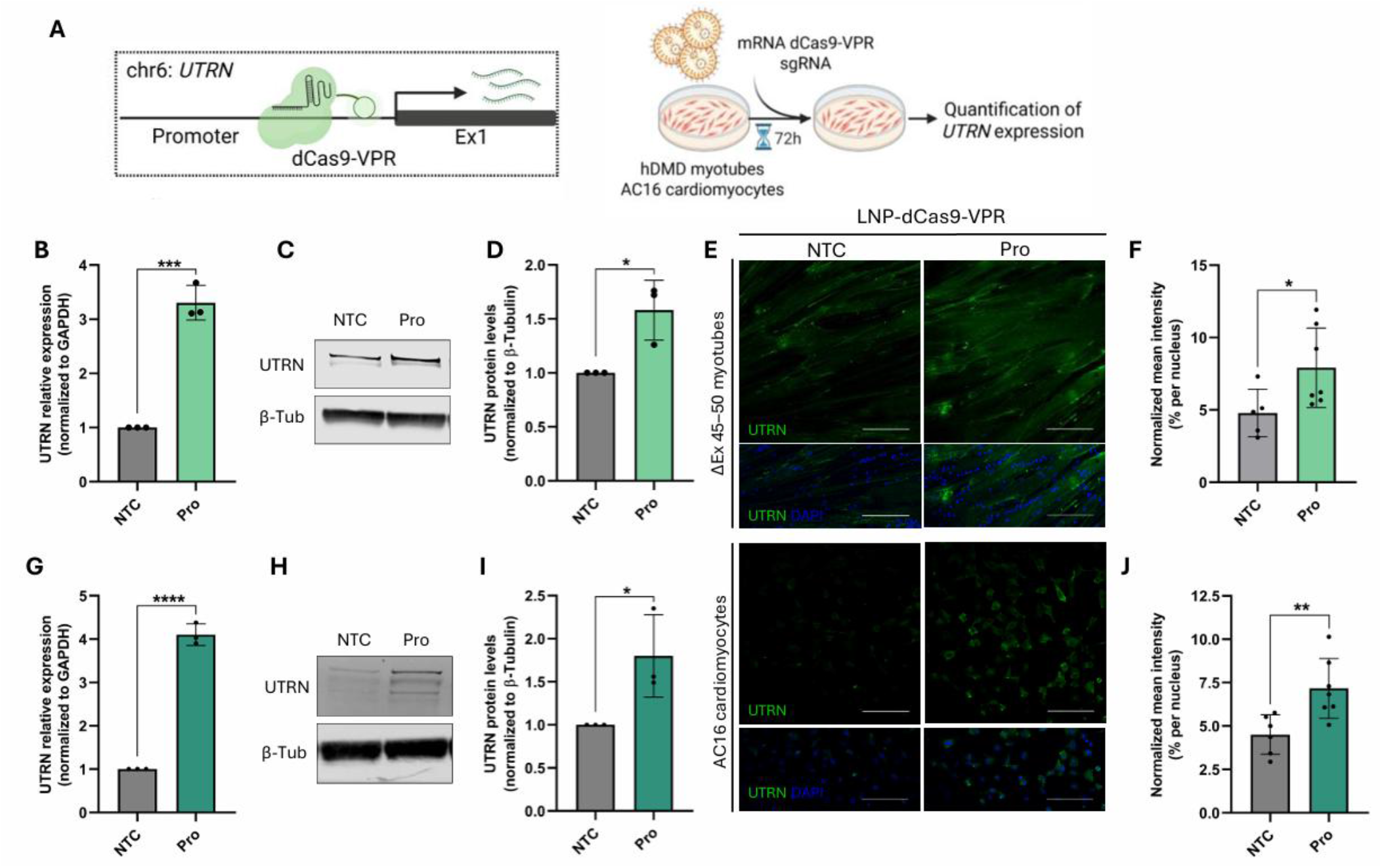
LNP-dCas9-VPR upregulates UTRN in ΔEx45–50 DMD myotubes and cardiomyocytes. (**A**) Schematic representation of utrophin activation strategy on the left and of the experimental workflow on the right. (**B, G**) RT–qPCR analysis of UTRN expression levels in ΔEx45–50 DMD myotubes (B) and AC16 cardiomyocytes (G); fold change is shown relative to the non-targeting control (NTC). (**C–D, H–I**) Western blot analysis of UTRN expression (C, H) and corresponding quantification (D, I) in ΔEx45–50 DMD myotubes (C–D) and AC16 cardiomyocytes (H–I), with β-tubulin as a loading control. Fold change is shown relative to NTC. (**E, J–F**) Immunofluorescence staining of UTRN (green) and nuclei (DAPI) in ΔEx45–50 DMD myotubes and AC16 cardiomyocytes (E), and corresponding quantification of UTRN fluorescence intensity normalized to the number of nuclei (F, J; n = 5–7 images per condition). Scale bar: 150 μm. NTC = sgRNA non-targeting control. Pro = sgRNA targeting the UTRN promoter. Data represent mean ± S.D. (n ≥ 3 biological replicates). Statistical analysis was performed using a two-tailed Student’s t-test. * p < 0.05, ** p < 0.01, *** p < 0.001, **** p < 0.0001. Each dot represents one sample.

Collectively, these results show that a single LNP formulation can efficiently deliver RNA cargos from short ASOs (∼20 nt) to large CRISPR-based effectors (∼6.7 kb) without re-optimization. This enables both genome editing and transcriptional modulation in multiple human muscle models, including post-mitotic systems that remain challenging for conventional transfection approaches.

## DISCUSSION

A major obstacle in the development and evaluation of RNA therapeutics for Duchenne muscular dystrophy is the lack of efficient delivery methods in differentiated human muscle cells. Post-mitotic myotubes represent one of the most physiologically relevant *in vitro* model for studying muscle biology and disease, as they recapitulate key features of mature muscle fibers, including multinucleation and expression of muscle-specific proteins [10]. However, their post-mitotic state and limited susceptibility to conventional transfection approaches have restricted their routine use in therapeutic development pipelines.

In this study, we demonstrate that a single commercially available lipid nanoparticle formulation enables robust RNA delivery directly into post-mitotic differentiated human myotubes and iPSC-derived myotubes, while remaining effective across additional human muscle models, including proliferating myoblasts, cardiomyocytes, and engineered 3D skeletal muscle tissues. Furthermore, we show that this platform supports the delivery of a broad range of therapeutic RNA cargos, from splice-switching antisense oligonucleotides to complex CRISPR-based genome editing systems.

Because genome editing outcomes are highly dependent on cellular context and cell-cycle status [11], approaches validated in proliferating myoblasts may not accurately predict delivery efficiency, editing outcomes, or therapeutic efficacy in mature muscle tissue. By enabling efficient RNA delivery directly in post-mitotic muscle cells, the platform described here provides a practical framework for evaluating emerging RNA therapeutics in physiologically relevant human models.

Here, we used a commercially available LNP formulation produced using a microfluidic mixer [32], yielding particles with homogeneous size and structure. This structural uniformity facilitates reproducible comparisons across RNA cargos and experimental models, enabling standardized evaluation of delivery performance across laboratories. In addition, efficient RNA encapsulation minimizes material loss and supports the generation of concentrated formulations suitable for screening applications. We also identified culture conditions as an important determinant of delivery efficiency, with growth medium significantly enhancing transgene expression compared to differentiation medium. While the underlying mechanism was not directly investigated, differences in medium composition may influence LNP uptake and intracellular trafficking, potentially through modulation of endocytic pathways or receptor availability. In particular, LNP internalization is known to occur via regulated endocytic mechanisms and may involve low-density lipoprotein receptor (LDLr)–dependent pathways, consistent with our detection of LDLr expression in both myoblasts and myotubes [19,33].

Across all conditions tested, LNP-mediated delivery resulted in robust RNA transfer and transgene expression, outperforming standard transfection reagents without detectable short-term toxicity [24,34,35]. These differences likely reflect fundamental mechanistic distinctions between delivery systems. While lipoplexes rely largely on nonspecific electrostatic interactions, LNPs provide a structurally defined and protective environment for mRNA encapsulation, thereby reducing extracellular degradation [36]. Moreover, the presence of ionizable lipids enables pH-dependent endosomal escape, a critical step that remains a major bottleneck in RNA delivery [37,38].

A key finding of this study is that a single LNP formulation was able to support efficient RNA delivery across multiple human muscle models without the need for protocol re-optimization. Following successful delivery in differentiated myotubes, we extended this approach to proliferating myoblasts, cardiomyocytes, and engineered skeletal muscle tissues. The ability to use the same formulation across distinct cellular systems simplifies experimental workflows, facilitates direct comparison of therapeutic strategies, and provides a standardized framework for preclinical evaluation. From a practical perspective, this versatility represents an important advantage for laboratories seeking to rapidly assess emerging RNA therapeutics in disease-relevant human models.

We further extended these observations to human 3D engineered muscle tissues, where LNPs efficiently transfected organoids both during tissue formation and in differentiated constructs containing multinucleated myofibers. While LNPs have been successfully applied in various organoid systems – including retinal, epithelial, and neural organoids [39–41] – their application to skeletal muscle organoids has not been extensively explored. In this context, our work represents, to our knowledge, the first demonstration of LNP-mediated RNA delivery in 3D human skeletal muscle organoids both during tissue formation and in pre-formed constructs, although with reduced penetration in the latter case. This likely reflects the increased structural complexity of mature tissues, where dense cellular organization and extracellular matrix components such as Matrigel may limit nanoparticle diffusion and penetration. Additionally, further optimization of LNP formulations tailored for skeletal muscle organoids is expected to enhance delivery efficiency. Although not tested here, this platform may be applicable to more complex multicellular muscle organoids, which better recapitulate tissue architecture and cellular heterogeneity [42].

Beyond transgene expression, we demonstrate that a single LNP formulation can efficiently deliver a wide range of therapeutic RNA cargos, spanning more than 300-fold in size. This includes short, chemically modified ASOs, which induced exon skipping and dystrophin restoration in post-mitotic myotubes, as well as larger and more complex CRISPR-based systems. Delivery of Cas9 nucleases, base editors, and CRISPRa constructs enabled multiple therapeutic outcomes, including reframing, exon skipping, and utrophin upregulation. Furthermore, the system supported co-delivery of mRNA together with sgRNAs, underscoring its high cargo versatility. Notably we observed that repeated LNP administration could increase editing efficiency in myotubes, suggesting that repeated dosing may represent a practical strategy to improve editing outcomes in post-mitotic systems, where genome modification cannot be amplified through cell division. These results were achieved using one LNP formulation without the need for re-optimization across cargos or cell types, highlighting a key practical advantage over conventional delivery approaches. Importantly, for genome editing studies, the possibility of using post-mitotic myotubes is of primordial interest as editing outcomes-including NHEJ and HDR for DNA-targeting editors, as well as gene expression effects for transcriptional and epigenome editors-are known to be influenced by the cell-cycle stage [11].

The ability of LNPs to deliver different RNA cargos required for genome editing approaches provides a clear advantage over viral vector systems. Viral vector production is time-consuming and resource-intensive, and it is poorly suited for the parallel evaluation of multiple CRISPR strategies or RNA designs. In contrast, the transient expression of mRNA delivered via LNPs represents a key benefit for CRISPR-based applications, as it limits prolonged Cas9 activity, which has been associated with increased risks of off-target editing and on-target genotoxicity [43,44]. This is particularly important when targeting non-dividing tissues such as skeletal muscle, where the absence of cell division prevents dilution of vector genomes and may exacerbate cell stress associated with sustained DNA double-strand breaks and Cas9 expression [45,46]. In addition, AAV vectors present intrinsic cargo size limitations (∼4.7 kb), which often require the use of dual-vector strategies for the delivery of large genome-editing constructs, such as split Cas9 or base editors [47]. These approaches rely on the co-delivery and reconstitution of the editing machinery, which can reduce overall efficiency and increase variability. Finally, unintended integration of AAV sequences at sites of Cas9-induced DNA double-strand breaks has been reported, highlighting potential safety concerns associated with persistent DNA-based delivery strategies [48,49].

Several limitations of this study should be considered. While the present study focused on a commercially available reference formulation, future studies comparing alternative muscle-targeted LNP compositions may further improve delivery efficiency and tissue specificity. First, we did not directly quantify the proportion of empty versus RNA-loaded LNP particles, nor we did systematically evaluate the optimal mRNA:sgRNA ratio, two parameters that are likely to influence delivery efficiency and could be further refined to maximize performance in specific applications [50–52]. Second, this study was limited to *in vitro* models. While our results demonstrate robust delivery across multiple *in vitro* muscle systems, translating LNP-based approaches treatment of muscle diseases such as DMD remains a major challenge. Although the clinical potential of LNP-mediated delivery has been validated, including successful *in vivo* genome editing in humans using LNP-delivered CRISPR–Cas9 targeting the TTR gene [20], current LNP formulations preferentially accumulate in the liver after intravenous injection and fail to achieve therapeutically relevant delivery to skeletal muscle [53]. While intramuscular administration has shown promising results [23,54], effective systemic targeting of skeletal muscle remains a key barrier. Overcoming this limitation will require further optimization of LNP composition and formulation parameters, which are known to critically influence cargo loading, stability and intracellular trafficking [55,56]. Strategies such as surface ligand conjugation have been explored to enhance tissue specificity, although their applicability to skeletal muscle remains to be fully established [57–59].

In the future, this LNP platform could be further explored to deliver CRISPR-based tools into autologous iPSC-derived myogenic cells, supporting the development of cell-based therapeutic strategies [60,61]. More broadly, the extensive cargo flexibility demonstrated here suggests that this LNP platform may be adapted to additional therapeutic modalities, including protein-based delivery systems such as ribonucleoprotein complexes, as well as small molecules [62,63].

Taken together, these findings provide, to our knowledge, the first systematic comparison of RNA delivery efficiency across both 2D and 3D myogenic models using a single LNP platform. By enabling efficient delivery of cargos ranging from ASOs to large CRISPR-based editors in differentiated myotubes, cardiomyocytes and engineered skeletal muscle tissues, this approach provides a practical framework for the rapid preclinical evaluation of emerging RNA therapeutics. Beyond DMD, the platform described here may facilitate the development and optimization of genetic therapies for a broad range of neuromuscular disorders.

## MATERIALS AND METHODS

### Cell culture

Immortalized healthy donor and patient-derived myoblasts (WT, DMD ΔEx51, ΔEx52, and ΔEx45–50) were generated from muscle biopsies and obtained from the Myobank-AFM cell repository at the Institute of Myology (Paris, France). Cells were cultured in skeletal muscle cell growth medium (C-23060, PromoCell) supplemented with 20% fetal bovine serum (FBS) and 1% penicillin–streptomycin (Invitrogen) and passaged using TrypLE Express (Gibco). For cell differentiation, myoblasts were seeded on collagen-coated plates (Corning) and allowed to reach ∼90% confluence. Differentiation into myotubes was induced by serum deprivation, achieved by switching to skeletal muscle cell differentiation medium (C-23061, PromoCell). The differentiation medium was refreshed every 2 days, and full differentiation into myotubes was reached within 3–5 days.

Human iPSC lines (WT and DMD ΔEx45) were originally generated from skin fibroblasts obtained from the Coriell Institute and provided by the Marseille Stem Cell Platform (Marseille Medical Genetics, France). These lines were previously engineered to stably express doxycycline-inducible MyoD and BAF60c transgenes [16]. Human iPSCs were maintained in mTeSR Plus medium (STEMCELL Technologies) and passaged using ReLeSR (STEMCELL Technologies) on matrigel-coated plates (Corning). Myogenic differentiation was performed as previously described [16]. Briefly, iPSCs were induced with doxycycline (200 ng/mL) for 24 hours to activate MyoD and BAF60c expression. The following day, iPSC colonies were detached, dissociated into single cells, and seeded at a density of 65,000 cells/cm² on Matrigel-coated plates (Corning) in growth medium (SKM02, AMSBIO). After two days, the medium was replaced with differentiation medium (SKM03, AMSBIO) supplemented with doxycycline (200 ng/mL). Differentiation medium was refreshed every 1–2 days, and full myogenic differentiation into iMyotubes was typically achieved within 3–4 days.

The AC16 cells (ATCC) were maintained in growth medium (DMEM/Nutrient Mixture F-12; Biowest, L0092) supplemented with 12.5% FBS (Thermo Fisher Scientific, 10270-106) and 1% penicillin–streptomycin (Invitrogen) and passaged using TrypleE (Gibco).

All cells were maintained at 37°C in a humidified incubator with 5% CO_2_. Cell lines were routinely tested for mycoplasma infection with PlasmoTest Mycoplasma Detection Kit (Invivogen).

### Generation of human 3D engineered skeletal muscles

3D human skeletal muscles were generated following established protocols [42,64]. Briefly, 10^6^ human immortalized myoblasts were resuspended in 120 μL of a matrix consisting of 3.5 mg/ml human fibrinogen (F3879, Sigma-Aldrich), 10% Matrigel (BD, 356230) and 3 U/ml thrombin (T7326, Sigma-Aldrich). The cell–matrix suspension was cast into agarose molds and polymerized for 2 h at 37 °C between paired polydimethylsiloxane (PDMS) posts (Dinabios/EHT Technologies, GmbH Hamburg). Following polymerization, constructs were maintained at 37°C in 5% CO₂ in growth medium (C-23060, PromoCell) supplemented with 33 μg/μL aprotinin (A3428, Sigma-Aldrich) to prevent fibrin degradation. After 48 h, 3D muscles were transferred to differentiation medium (C-23061, PromoCell), also supplemented with aprotinin, to induce myogenic differentiation. The medium was replaced every other day.

### *In vitro* transcription (IVT) of mRNA

Plasmids encoding the genome editors under a T7 promoter (Addgene, plasmid #185910; plasmid #63798) were linearized downstream of the stop codon and purified using the QIAquick PCR Purification Kit (QIAGEN). One microgram of linearized plasmid was used as template for IVT with the MEGAscript™ T7 Transcription Kit (Ambion), including co-transcriptional capping with CleanCap AG (TriLink Biotechnologies). The IVT reaction was performed according to manufacturer’s instructions with the following modification: the GTP concentration was reduced to 3.0 mM, and CleanCap AG was added to a final concentration of 12.0 mM, resulting in a 4:1 Cap:GTP ratio, as previously described to enhance capping efficiency [65]. After transcription, the reaction was treated with DNase I to remove residual DNA template. Polyadenylation was performed using the Poly(A) Tailing Kit (Invitrogen) according to the manufacturer’s protocol. The resulting mRNA was precipitated with lithium chloride, resuspended in TE buffer, aliquoted, and stored at -80°C. mRNA size and integrity were assessed by electrophoresis on 1% agarose gel.

### Synthetic guide RNA and ASO

Chemically synthesized sgRNAs were purchased from either Synthego or Integrated DNA Technologies (IDT). The antisense oligonucleotide (ASO) was synthesized as a fully modified 2′-O-methyl oligonucleotide and obtained from IDT. All sgRNAs and ASOs were aliquoted upon receipt and stored at -80 °C until use. The sequences are provided in the supplementary material.

### LNP formulation and characterization

LNPs were formulated by mixing GenVoy-ILM lipids (Cytiva) with RNA cargo at an N/P molar ratio of 6:1 using the NanoAssembl Spark microfluidic system (Cytiva), following the manufacturer’s instructions. For genome-editors formulations, Cas9-based mRNA editor and sgRNA were mixed at equal mass in the aqueous phase prior to encapsulation. GFP mRNA cargo was ordered from GenScript (eGFP mRNA Cap1, 5-MOU) and Cas9 nuclease mRNA cargo was ordered from TebuBio (cleancap Cas9 mRNA, L-7606-100). Immediately after formulation, LNPs were diluted 1:1 in PBS. Total RNA concentration and encapsulation efficiency were quantified using the Quant-iT™ RiboGreen assay (Invitrogen), according to the manufacturer’s protocol. Because the ASOs carry fully modified 2′-O-methyl backbone chemistry, they cannot be quantified accurately using the RiboGreen fluorescence assay. As the microfluidic mixing step results in negligible volume loss, the ASO concentration in the final LNP formulation was therefore calculated based on the initial input amount and the final dilution factor. LNP size and polydispersity were measured using a NanoSight NS300 instrument (Malvern Panalytical). Three 90-second videos were recorded at 20 °C using the 488 nm laser module. Formulated LNPs were stored at 4°C and used within 7 days for transfection experiments.

### LNP transfection

To define the optimal media for myotubes transfection, patient-derived myotubes were transfected with LNP-GFP mRNA in growth media or differentiation media supplemented with 0.2 µg/mL of ApoE3 (Thermo Fisher Scientific), 0.2 µg/mL of ApoE4 (Thermo Fisher Scientific), or 20% FBS. For dose optimization experiments with LNP-GFP mRNA, 50,000 immortalized patient-derived myoblasts and 40,000 AC16 cardiomyocytes were seeded into 24-well plates, aiming to reach approximately 50% confluency the following day. LNP-GFP formulations were added directly to the respective cell culture medium at the indicated doses (see figure legends). 24 hours later, GFP-positive cells were quantified by flow cytometry (Beckman Coulter), and fluorescence images were acquired using an EVOS epifluorescence microscope (Thermo Fisher Scientific). For optimization experiments in differentiated cells, immortalized patient-derived myoblasts or iPSCs were first differentiated into myotubes in 24-well plates. Once differentiation was complete, differentiation medium was replaced by serum-containing medium (Myotubes: skeletal muscle cell growth medium supplemented with 20% FBS and 1% penicillin–streptomycin; iMyotubes: SKM03 medium with 5% FBS and 200 ng/µL doxycycline) to enhance LNP internalization. LNP-GFP formulations were then added directly to this medium at the indicated doses (see figure legends). GFP fluorescence was assessed the following day using a fluorescent plate reader, and images were acquired with the EVOS microscope.

For delivery of RNA therapeutics in differentiated cells, immortalized patient-derived myoblasts or iPSCs were first differentiated into myotubes in 12-well plates. Upon completion of differentiation, differentiation medium was replaced by serum-containing medium as described above. Unless otherwise specified, LNPs containing 4 µg of encapsulated mRNA were added in serum containing medium. The following day, the medium was replaced with differentiation medium. Three days after LNP treatment, cells were harvested for DNA, RNA, and protein extraction to evaluate therapeutic efficacy.

### Lipofectamine-based transfection

A total of 50,000 immortalized patient-derived myoblasts and 40,000 AC16 cardiomyocytes were seeded in 24-well plates, aiming to reach approximately 50% confluency the following day. For transfection, 500 ng of GFP mRNA was mixed with 1.5 µL of Lipofectamine 3000 (Thermo Fisher Scientific) and applied to the cells according to the manufacturer’s instructions. GFP expression was quantified 24 hours post-transfection by flow cytometry.

### Nucleofection

A total of 150,000 myoblasts were nucleofected with 500–2000 ng of GFP mRNA or with 2,000 ng of ABE8e IVT mRNA together with 60 pmol of synthetic sgRNA, using the P5 Primary Cell buffer (Lonza) and the C2C12 program on the 4D-Nucleofector system (Lonza). GFP expression was quantified 24 hours post-nucleofection by flow cytometry. For base-editing experiments, the culture medium was replaced the day after nucleofection, and cells were harvested 3 days post-nucleofection for DNA extraction and assessment of editing efficiency.

### Flow Cytometry analysis

24 hours after LNP-GFP treatment, immortalized patient derived myoblasts or AC16 cardiomyocytes were washed with 1× PBS (Gibco) and dissociated using TrypLE Express (Gibco). TrypLE was inactivated, and cells were washed and resuspended in PBS prior to analysis. For LDLr quantification, 250,000 immortalized patient-derived myoblasts or AC16 cardiomyocytes were incubated for 20 min with 1 µL of LDLr antibody (1/200, clone 301, Invitrogen). Flow cytometry was performed using a CytoFLEX S cytometer (Beckman Coulter). After gating for live single cells (doublet exclusion), approximately 10,000 positive events per sample were collected to quantify percentage of positive cells and mean fluorescence intensity For LDLr quantification, data were analysed using CytExpert software (Beckman Coulter). For viability assessment, cells were incubated with 7-Aminoactinomycin D (7-AAD, Thermo Fisher) for 15 minutes at room temperature before acquisition. Unstained cells were included as negative controls.

### Evaluation of GFP intensity into myotubes

One day after LNP-GFP transfection of myotubes or iMyotubes, the culture medium was removed and replaced with PBS. GFP fluorescence was measured immediately using an EnSpire plate reader (PerkinElmer) with an excitation wavelength of 485 nm and an emission wavelength of 520 nm. GFP intensity was reported in arbitrary units after subtraction of the background signal (PBS only).

### DNA extraction, PCR and Sanger sequencing

Genomic DNA was extracted from frozen cell pellets with QIAamp DNA Micro Kit (Qiagen). DNA purity and concentration were assessed with a Nanodrop spectrophotometer. For each reaction, 50 ng of genomic DNA were used as template for PCR amplification with KAPA2G Fast ReadyMix (Kapa Biosystems, Wilmington, MA, USA). Primer sequences are listed in the supplementary material. PCR products were loaded on a 1% agarose gel to confirm amplicon size and purity before submission for Sanger sequencing (Genewiz company, Takeley, UK). Editing efficiency was further assessed using chromatogram-based analysis tools: TIDE [66] for CRISPR nuclease–mediated editing and EditR [67] for base-editing quantification.

### RNA extraction, reverse transcription, quantitative PCR, and PCR

Total RNA was isolated using the RNeasy Micro Kit (Qiagen, Hilden, Germany). RNA purity and concentration were assessed by Nanodrop, and 500 ng of RNA was used for reverse transcription with the Transcription First Strand cDNA Synthesis Kit (Roche, Basel, Switzerland). For gene expression analysis, quantitative PCR was performed using Maxima SYBR Green/ROX Master Mix (Thermo Fisher Scientific, Waltham, MA, USA) on a LightCycler 480 system (Roche, Basel, Switzerland). Relative gene expression levels were calculated using the 2^−ΔΔ^*^Ct^* method, with *GAPDH* serving as the housekeeping gene for normalization. For exon 44 and exon 50 skipping analysis, PCR on cDNA was carried out using KAPA2G Fast ReadyMix (Kapa Biosystems, Wilmington, MA, USA). PCR products were separated on 1% agarose gels containing SYBR Safe DNA stain (Invitrogen, Carlsbad, CA, USA) and band intensities were quantified using a UV imaging system. Bands corresponding to the expected exon-skipped products were excised and purified using the NucleoSpin Gel and PCR Clean-up Kit (Macherey-Nagel, France). Splice junctions were confirmed by direct Sanger sequencing (Genewiz company, Takeley, UK). Primer sequences are listed in the supplementary material.

### Protein extraction and Western Blot

Cells were lysed in RIPA buffer (R0278, Sigma-Aldrich) supplemented with a protease inhibitor cocktail (P8340, Sigma-Aldrich). Following BCA protein quantification using the Pierce™ BCA Protein Assay Kit (ThermoFisher), 10–20 μg of total protein was mixed with NuPAGE™ LDS sample buffer (NP0007, Invitrogen) and β-mercaptoethanol (11528926, Fisher Scientific). Samples were heat-denatured for 5 min at 100°C and loaded onto NuPAGE 3–8% Tris-Acetate Midi Gels (Novex, Life Technologies), then transferred onto PVDF membranes (Millipore). Membranes were blocked for 1 h at room temperature in Odyssey blocking buffer (926-41090, LI-COR) and incubated overnight at 4°C with the following primary antibodies: mouse anti-dystrophin (1:50, NCL-DYSB, Leica), mouse anti-utrophin (1:100, clone 84A, SC-33700, Santa Cruz Biotechnology), and rabbit anti-beta Tubulin (1:2000, ab6046, Abcam). After washing, membranes were incubated for 1h at room temperature (RT) in the dark with the following secondary antibodies: IRDye 680RD goat anti-rabbit (LI-COR) and IRDye 800CW donkey anti-mouse (LI-COR). Fluorescent signals were detected using the Odyssey Imaging System (LI-COR Biosciences). Band intensities were quantified using ImageJ, and target protein levels were normalized to beta tubulin.

### Processing of 3D engineered skeletal muscles

3D muscles were fixed overnight at 4 °C in PBS containing 4% PFA (JI9943.K2, Thermo Scientific). After fixation, samples were equilibrated in 30% sucrose (S0389, Sigma) for 48 h at 4 °C. The muscles were then embedded in PBS supplemented with 7.5% gelatin (G-1890, Sigma) and 10% sucrose, and snap-frozen in isopentane at –50 °C. Transverse sections (10 μm) were obtained using a Leica cryostat and mounted onto glass slides.

### Immunostaining of cells and 3D engineered skeletal muscles

For immunostaining of AC16 cardiomyocytes and myotubes, cells were seeded in 35 mm cell culture microscopy dish (µ-Dish^35mm, high^, Ibidi). After LNP transfection, cells were washed with PBS and fixed in 4% PFA for 15 min at RT. Following PBS washes, cells were permeabilized with 0.25% Triton X-100 for 5 min and blocked in 10% FBS for 45 min at RT. Cells were then incubated for 2 h at RT or overnight at 4°C with the following primary antibodies: mouse anti-dystrophin (1:50, NCL-DYSB, Leica), mouse anti-utrophin (1:100, clone 84A, SC-33700, Santa Cruz Biotechnology); GFP Polyclonal Antibody, Alexa Fluor™ 488 (1:500, Invitrogen, A-21311). For non-conjugated antibodies, after PBS washes cells were incubated for 1-2 h at RT in the dark with a goat anti-mouse alexa 594 conjugated secondary antibody (1:500, A-11005, Life Technologies). Cells were washed and mounted using DAPI Fluoromount-G (00-4959-52, Invitrogen). Images were acquired on a LEICA SP8 Confocal Microscope (Zeiss, Germany) using a HC PL APO CS2 20X / 0.75NA dry objective (Zeiss, Germany). For quantification of UTRN expression, images were processed using a custom-made script on Fiji Plateform [68]. The Dapi image was segmented after applying a Gaussian Blur filter (sigma=2), using the automatic minError threshold [69]. As the nuclei were not clearly segmented, we iterated the method using once again a Gaussian Blur filter (sigma=7), and then using the automatic Max Entropy threshold [70]. Finally, we used the watershed build-in algorithm, and the Analyze Particle build-in Fiji function to identify and count the number of nuclei per image. For each image, the mean intensity of the channel of interest was computed on the whole image, and the intensity was normalized by the number of nuclei, resulting in an intensity score for each image: intensity score = Mean Intensity / number of nuclei * 100.

### Statistical Analysis

Statistical analyses were performed using GraphPad Prism 7. Significance was assessed using two-tailed Student’s *t*-tests unless otherwise specified; additional statistical tests are indicated in the corresponding figure legends. Differences were considered significant at the following thresholds: *p* < 0.05 (\**), p* < 0.01 (\*\**), p* < 0.001 (***), and *p* < 0.0001 (****). Data are presented as mean ± S.D., and *n* refers to the number of independent biological replicates included in each comparison.

## ACKNOWLEDGMENTS

We acknowledge Genethon “Imaging and Cytometry Core Facility”, especially Peggy Sanatine, Simon Jimenez and Jérémie Cosette for technical support in confocal acquisition and images analysis, and “Histology platform”, especially Fanny Bordier, for providing histological section of 3D skeletal muscle tissues. We gratefully acknowledge Aurélien Jacob and Mallaury Laverry for their valuable technical advice, for critical reading of the manuscript and for sharing some RNA and LNP reagents. We acknowledge Pr. Philippe Menasché and Valerie Bellamy for providing the AC16 cell line, as well as Annarita Miccio and Simone Amistadi for sharing the plasmid encoding for dCas9-VPR. We gratefully acknowledge the, Ile-de-France Region, Conseil Departmental de l’Essonne (ASTRE), GIP, Genopole and GenoTher for the financial help to purchase equipment. We thank the whole Amendola’s laboratory for fruitful discussion. This work was funded by the European Union (Horizon Europe project no. 101080690 – MAGIC), the UK Research and Innovation (UKRI) under the UK government’s Horizon Europe funding guarantee grants no. 10080927, 10079726, 10082354, 10078461, the Swiss State Secretariat for Education, Research and Innovation (SERI) (www.magic-horizon.eu; MA), grants from the Agence Nationale de la Recherche (HemoLen ANR-20-CE17-0016-0 (MA), PEMGeT ANR-22-CE17-0028-02 (MA); IRIS ANR-21-CE14-0063-03 (MA); NEEDED ANR-24-CE18-3712-01 (MA), GenoTher ANR-23-BIOC-0003 (MA)), AFM-telethon (MA), INSERM (MA) and University of Évry Val d’Essonne (MA, PG). The authors also acknowledge funding from the European Union (EDITSCD grant 101057659, https://editscd.eu/ (MA); ERDERA grant 101156595 (MA); COST action grant [GeneHumdi-CA21113] (MA). Views and opinions expressed are however those of the author(s) only and do not necessarily reflect those of the European Union or HaDEA. Neither the European Union nor the granting authority can be held responsible for them.

## AUTHOR CONTRIBUTIONS

P.G., D.L. and M.A conceived the study. P.G., D.L. M.M., R.K., M.R., G.B., G.S. contributed to the execution and/or analysis of the experiments; K.M. provided cell lines; F.S.T., and S.A. supervised the organoid, the LNP characterization, and the iPSC experiments, respectively. M.A. supervised the study. P.G., D.L. and M.A. wrote the manuscript.

## DECLARATION IF INTERESTS

The authors declare no competing interests.

### Declaration of generative AI and AI-assisted technologies in the writing process

During the preparation of this work, the authors used generative AI tools to improve clarity and streamline the text to meet the journal’s space constraints. All scientific content, data interpretation and conclusions were critically reviewed and edited by the authors, who take full responsibility for the integrity and accuracy of the published work.

## SUPPLEMENTARY INFORMATION

### Supplementary material

ASO, sgRNAs, and primers used in this study.

